# Preserving predictive information under biologically plausible compression

**DOI:** 10.1101/2025.03.12.642864

**Authors:** Sylvia C. L. Durian, Kyle Bojanek, Olivier Marre, Stephanie E. Palmer

## Abstract

Retinal ganglion cells (RGCs) show high convergence onto their downstream projections, which poses a problem for information transfer: how can information be preserved through a synaptic layer that has significantly more inputs than outputs? Lossy compression suggests many efficient, yet computation-agnostic, methods for reading out input stimuli or activity patterns. Focusing on prediction as a ubiquitous computation in the brain, we compare compressions that explicitly retain predictive information to common neural compression frameworks that do not. We find evidence that downstream areas may compress their retinal inputs in a way that allows them to perform optimal predictive computations across many natural scenes. Other sensory systems also exhibit compression in their processing hierarchies, such as at the glomeruli stage in the olfactory system, and we hope that our framework will be useful in cases where it is not yet known how information about a specific computation is maintained under compression.

**SIGNIFICANCE STATEMENT:** Producing successful behavior, such as escaping predators, requires the visual system to overcome significant sensory processing delays by predicting the future state of the world. Neurons in the eye capture some of the most predictive features of visual information, but this information must be accessible to downstream areas that receive synaptically compressed inputs from the retina. We tested how carefully retinal activity must be compressed to preserve predictive information in natural scenes. Biologically plausible compressions that are agnostic to prediction can preserve substantial future information, but only compression optimized for this task extracts generalizable motifs that allow it to predict in any natural scene. This suggests that downstream circuits may optimally compress their inputs specifically for the task of prediction.

## INTRODUCTION

Several sensory pathways exhibit both compression and expansion at different computational stages, as shown in neural recordings from successive regions in a processing hierarchy [1–3], and to help computational flow through layers of deep neural networks [4, 5]. Compression in particular poses a problem for information transfer, since it makes it difficult to retain all the input information. Previous work has used a compressed-sensing framework [6–8] to suggest that input stimuli or input neural activity patterns can be preserved under compression using random projections [9, 10], which was validated in an idealized network [11, 12]. However, the job of neural systems is to drive downstream behavior, not to reconstruct inputs, and not all input features are important for behavior. Therefore the efficient way to *coarse-grain*, or compress, would be to keep only the bits that matter for behavior. We explore this possibility using an information bottleneck approach [13–15], which is an established framework for studying compression achieved by throwing away task-irrelevant information.

Retinal ganglion cells (RGCs), the output cells of the retina, are particularly suited to studying how information about biologically relevant computations can be preserved under coarse-graining. RGC axons form the optic nerve, the sole channel carrying information about the visual world to the brain [16], which then synaptically converges onto several retinal projection areas. For instance, RGCs provide converging inputs to the lateral geniculate nucleus of the thalamus at an average rate of 10-1 in mice [17], although the number of inputs that significantly affect postsynaptic activity may be closer to 3 [17, 18]. Another key projection area is the optic tectum (OT in amphibians, fish, reptiles, etc.; superior colliculus in mammals), which receives input at a rate of roughly 6-1 in mice [19, 20].

Moreover, a large body of previous work has identified many important computations in the retina which downstream areas must be able to read out from their compressed inputs. For example, RGCs show a variety of nonlinear processing characteristics associated with motion prediction [21–24], which is necessary to avoid predators, catch prey, and navigate complex environments. In contrast to theories of predictive coding in which differences from an expected signal are encoded to efficiently transfer information [25–27], here we study bottom-up encoding of the predictive features of spatiotemporal structure in sensory stimuli. RGC population activity has internal temporal correlations which provide a near-optimal representation of future information about the external sensory world [15], indicating that prediction is an essential computation in the retina. The OT has a central role in making rapid eye and head movements in response to visual stimuli [28, 29], and is involved in fast innate behaviors like escaping and freezing in mice and amphibians such as salamanders and toads [30–33], suggesting that it relies on predictive information from the retina in order to drive successful behavior. We aim to understand how downstream areas like OT are able to perform predictive computations using the compressed information they receive from RGCs.

Information bottleneck theory and the animal’s behavioral goals guide us to hypothesize that downstream areas carry a near-optimal but compressed representation of the future of their natural environment. We compare optimal and prediction-agnostic coarse-grainings of RGC inputs to the full, uncompressed predictive information to determine how much information can be preserved under compression without taking readout of the most likely future input into account. If a prediction-agnostic coarse-graining can reach near-optimal readout, downstream areas might not waste energy and resources to actively optimize their RGC inputs. However, regardless of how compression is achieved, the amount of information received by a downstream neuron depends on which RGCs it receives input from. Downstream circuits decode the visual scene from action potentials generated by a diversity of RGC types [34, 35], which encode different visual features [36]. The retina’s output channels relay these different features to the brain, but our understanding of this process is complicated by the fact that some RGC types simultaneously code for multiple features in species such as mice and salamanders [37–39]. This can be beneficial: activity from across multiple direction-selective ganglion cell (DSGC) subtypes in mice enhances decoding of visual features and can help resolve ambiguities in single-cell responses [38, 40, 41]. The direction-selectivity of DSGCs is largely attributed to GABAergic input from starburst amacrine cells [38, 42], illustrating how differing inputs to RGCs lead to their encoding of different visual features. This motivates us to explore how RGC input composition affects coarse-grained outputs. We hypothesize that downstream areas like OT coarse-grain together their convergent inputs to optimally preserve predictive information in natural scenes.

We use methods from information theory to investigate optimal downstream readout of predictive information and show that common neural coarse-graining functions that do not explicitly take the future into account can preserve a surprisingly high proportion of the optimal predictive information. However, an efficient coarsegraining should ideally be able to predict the future in any natural scene [43–45], and we demonstrate that only a coarse-graining function explicitly optimized for prediction can also retain near-optimal predictive information in across scenes. This generalization suggests that downstream mechanisms may explicitly optimize their compressed RGC inputs for prediction. Finally, efficient compression depends not only on the coarse-graining algorithm but also on which cells are combined, and we determine that coarse-graining inputs from RGCs with shared bipolar inputs results in significantly more predictive information than combining random RGCs. Our results suggest a biologically plausible way for downstream areas like OT to efficiently encode the compressed predictive information they receive from the retina.

## RESULTS

We use dense extracellular recordings from larval tiger salamander RGCs that were made while presenting the retina with naturalistic movies [46](Fig. 1A). In this dataset, several movies were chosen to represent the wide variety of scenes that a salamander might encounter during its lifetime (Fig. 1B). These include trees moving in the wind, grasses rustling, fish swimming, and woodland underbrush as viewed by a moving camera to imitate the animal’s self-motion. Movies were presented in pseudorandom order and a white noise checkerboard stimulus was presented at the beginning and end to probe the classical receptive fields. Salamanders are an intriguing model species because they hatch underwater and move onto land as juveniles, then alternate between aquatic and terrestrial lifestyles as adults [44, 45]. However, their retinal structure remains largely the same across these stages [45, 47]. This allows us to study a wide diversity of natural scenes using larval population responses.

**FIG. 1.**
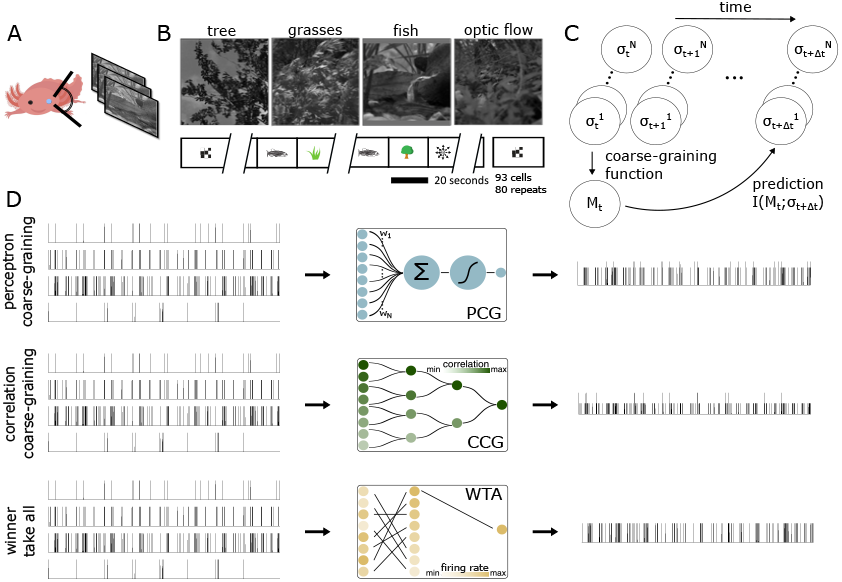
Behavioral paradigm and model. A) Simultaneous extracellular recordings were made from the ganglion cell layer of a larval tiger salamander while stimulating the retina with natural movies. B) Example natural movie frames. The optic flow movie is taken by a moving camera traveling through undergrowth. C) The activity of *N* neurons is coarse-grained down to one meta neuron *M* (*t*), which is used to read out future information. D) Schematics of our coarse-graining functions. The final traces represent the firing probability of the meta neurons over time. Top: the perceptron coarse-graining (PCG) is a weighted sum of inputs passed through a sigmoid nonlinearity. Middle: correlation coarse-graining (CCG) greedily pairs inputs based on their pairwise correlations, then successively sums and normalizes them. Bottom: winner take all (WTA) outputs the spike train of the input cell with the highest firing rate.

We represent the activity of a group of *N* neurons as a binary pattern 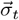, where 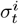 represents the activity of cell *i* at time *t* with a 1 if there was a spike and 0 if there was not. To study downstream computations, we coarse-grain *N* cells into one *meta neuron, M*_*t*_, which has 2 response states that depend probabilistically on the input activity 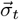. We then test how well the meta neuron retains information about future input activity (Fig. 1C):

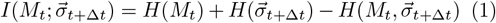

where *H* denotes entropy. While the goal of predictive mechanisms is to predict the future state of the external world, we measure predictive information this way because retinal projection areas do not have direct access to stimulus information, only the information in their inputs. However, in a simple stimulus, information about future input activity is related to information about the future of the stimulus, and optimal readouts of one are likely to be near-optimal readouts of the other [48]. We calculate the joint and marginal entropies using the joint probability, which is convenient to write as

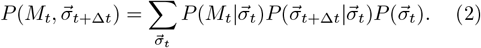

### Computation-agnostic compression preserves considerable predictive information

To test how we can extract compressed predictive information in practice, we compare coarse-grainings that are agnostic to the future to an optimal predictive coarsegraining. We model an optimal downstream neuron with a single-layer perceptron coarse-graining (PCG) [49], which is a linear weighting of inputs followed by a sigmoid nonlinearity (Fig. 1D top). This is a biologically feasible, single-step computation. The inputs to the PCG represent inputs from the dendrites, the weights represent synapses, and the output represents axonal output. The output of the coarse-graining is the conditional probability that the meta neuron spikes, given the input. Previous work has shown that PCG preserves reasonable predictive information in retinal data [15, 21], and that optimal PCG readouts are learnable via spike timing dependent plasticity (STDP) rules [48]. We compare PCG to a model downstream neuron that is biologically feasible but does not take the future into account, which we call correlation coarse-graining (CCG). Inputs are paired greedily based on their pairwise correlation, then summed and normalized at each step, again outputting a spiking probability (Fig. 1D middle). Meta neurons constructed in this same way from hippocampal populations have well-developed place fields [50], indicating that CCG can preserve biologically relevant information. We consider CCG as averaging with a Hebbian flavor [51], which we contrast with a Winner Take All (WTA) coarse-graining (Fig. 1D bottom), that is also prediction-agnostic. While we do not test every possible prediction-agnostic coarse-graining function, these two are both biologically realistic and are established computations used by the brain [52–54].

One key difference between the optimal and agnostic coarse-grainings is that the PCG weights are optimized for a particular computation, such as prediction, while CCG and WTA are not. We maximize predictive information in PCG while constraining its mean firing rate to match that of the corresponding CCG meta neuron to ensure that the output rate is biologically plausible. In fact, allowing the output firing rate of PCG to vastly exceed its mean input firing rate expands the dynamic range of PCG (Fig. 2A) in an unfair way by skirting around the coarsening constraint. We choose a small convergence rate of *N* = 4 cells to 1 meta neuron, since we are interested in modeling downstream areas involved in short-timescale prediction, such as OT, which has a similarly small convergence rate [19, 20]. With only 1 output meta neuron, future information is relatively constant as a function of the number of inputs *N* (Fig. 2C), which validates our choice of small *N*. The plateau in information suggests that a downstream neuron might not gain much predictive information by further local coarsening; instead, a small pool size may be optimal.

**FIG. 2.**
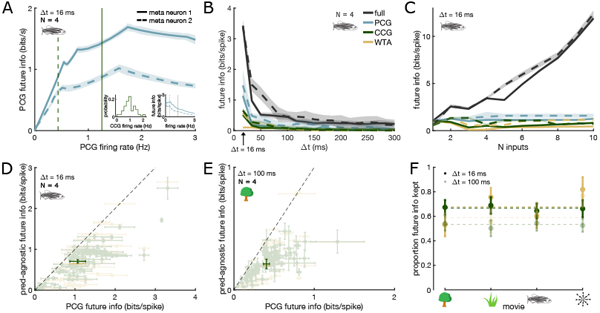
CCG and WTA preserve considerable predictive information compared to PCG. A) Future information maintained by PCG in the fish movie as a function of its firing rate in 2 example meta neurons. Both meta neurons were constructed by coarse-graining 4 input cells together. Error bars are standard deviations obtained by bootstrapping. Vertical lines denote firing rates of the corresponding CCG meta neurons constructed from the same inputs. Left inset: distribution of CCG firing rates in *n* = 73 meta neurons. Right inset: Future information in the two PCG meta neurons normalized by their firing rates. B) Future information maintained in the same 2 meta neurons (meta neuron 1, solid; meta neuron 2, dashed) at different prediction intervals. PCG is rate matched to CCG. C) Future information maintained by coarse-graining *N* input cells into 1 meta neuron, in 2 meta neurons of each input size (meta neuron 1, solid; meta neuron 2, dashed). D-E) Future information retained by CCG and WTA meta neurons (*n* = 73) compared to PCG meta neurons in different movies and at two prediction intervals. Each transparent point represents 1 meta neuron made from 4 random input cells. Opaque points are medians. Error bars are standard deviation obtained by bootstrapping. F) Proportion future information retained by CCG and WTA compared to PCG in each movie and at two prediction intervals, calculated from the data in D) and E) and summarized with the median. Error bars are standard error.

By design, PCG maintains more future information than the agnostic compressions (Fig. 2D-F) but surprisingly, both CCG and WTA retain a high amount of predictive information. This result is along the same lines as recent findings showing that random projections preserve input information in cortex [9, 12]. Importantly, we compared these coarse-grainings to optimal and find that they are both able to maintain a high proportion of the possible predictive information. This points to the possibility that downstream areas may not need to optimize their inputs in order to effectively use predictive information from RGCs to drive behavior.

### Optimal coarse-graining generalizes

Salamanders and other organisms that experience a wide variety of natural scenes during their lifetimes [43– 45] must be able to perform predictive computations in many environments. Much like orientation tuning of visual cortical neurons is invariant to changes in contrast [55, 56], efficient coding of predictive information should be scene invariant. Metabolic [57] and temporal constraints [58, 59] make it more efficient to use a coarsegraining algorithm which is flexible enough to successfully predict the future in any environment. While the explicit predictive power of PCG did not offer an overwhelming advantage over CCG and WTA in a given scene, it could help quickly adapt the system for prediction across a wide variety of scenes by extracting generalizable predictive encoding motifs. To investigate this, we train our three coarse-grainings on data pooled across scenes, then test them on an example movie (fish; Fig. 3B). This amounts to calculating 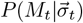 in Eq. 2 from the pooled training data and calculating 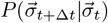 and 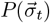 from the test data.

**FIG. 3.**
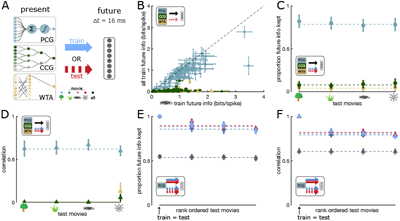
Predictive PCG generalizes across natural scenes. A) PCG, CCG, and WTA are trained and tested on different movies to investigate generalization. All data shown are for a prediction horizon of Δ*t* = 16 ms. B) We trained the three coarse-grainings trained on pooled data, then tested them on the fish movie (y-axis). This is compared to the amount of information that was retained by training and testing each coarse-graining on the fish movie. C) Mean proportion future information retained by PCG, CCG, and WTA trained on pooled movies, then tested on individual movies. Information from each type of coarse-graining is normalized by information in the test movie found by the *same* coarsegraining method. Errors bars are standard error and dashed lines denote means. D) Mean correlation coefficients between the coarse-grainings trained on pooled data and tested on different individual movies compared to the coarse-grainings trained and tested on the same movie. E) Proportion future information kept in the test movies by PCGs trained on different individual movies. Random PCGs (gray) are rate matched. X-axis is test movies sorted in descending order. F) Mean correlation coefficients between PCGs trained on different individual movies.

PCG meta neurons trained on the pooled data have comparable prediction performance to PCG meta neurons trained on the test movie, while CCG and WTA meta neurons trained on the pooled data fail to generalize to the test data. We quantify this in the proportion of future information kept by the coarse-grained meta neurons, which we normalize by the future information retained in the relevant test movie by the *corresponding* coarse-graining (Fig. 3C). CCG and WTA might seem more efficient because they are more simple, but they do not allow for scene invariant prediction, whereas PCG may be flexible enough to predict its next inputs in any natural scene. We test generalization in PCG more carefully by training on individual movies; this allows us to determine whether PCG can generalize to a movie it has never seen before, and to rule out the possibility that its high generalization performance in Fig. 3C was due to learning the similar scene statistics in the tree and grasses movies, for example, rather than learning something more general about natural scene statistics. We find that indeed, PCGs trained on individual movies retain a high proportion of future information in other movies, and more than rate-matched random PCGs. This indicates that retinal projection areas may be able to solve a generalizable prediction problem for natural scene statistics, if they can wire up converging downstream connections in a way that retains predictive information. Related work from our lab has shown that a bottom-up coarse-graining of the retina identifies the slow features of natural scenes as both the most important *and* most general across scenes [60]. Because physical objects have momentum, slow features are also the most predictable. Therefore, our result is a product of the retina’s selection of specific predictive features in the natural movies, as well as the fact that natural scenes have predictive-generalizable features because they share some common scene statistics.

### Cells with shared inputs maintain more predictive information

In practice, the amount of predictive information available to a downstream neuron depends on which RGCs it receives input from, regardless of the coarse-graining function used by downstream circuits. How do we find which groups of cells the brain might coarse-grain together? Classical efficient coding theories [61, 62] suggest that cells with overlapping information can be efficiently coarse-grained by reducing redundancy. This is easily understood in the case of CCG: performing CCG on two perfectly correlated cells would perfectly preserve the original activity. Cells that share input information (Fig. 4A) must automatically share some overlapping information in their responses, and it is possible that the brain exploits this redundancy by coarse-graining them together. We hypothesize that meta neurons constructed from RGCs with shared bipolar inputs carry more predictive information than ones constructed from random inputs.

**FIG. 4.**
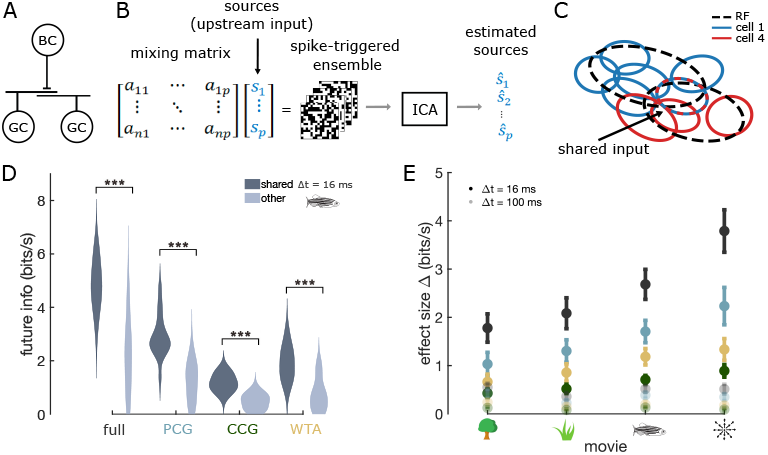
Cells with shared inputs carry more information about the future. A) Schematic of shared bipolar cell input to RGCs. B) Schematic of spike-triggered independent component analysis (ST-ICA) method used to model shared bipolar cell input. C) Spatial subunits recovered from ST-ICA analysis for two example cells who share a subunit. Dashed black lines represent the receptive fields (RFs) of the cells. D) Distributions of future information in meta neurons (and full neurons) constructed from cells sharing at least one common input, and meta neurons without any common input. E) Summary of effect size (difference in means between the two distributions) in each movie and at two prediction intervals.

We identify putative shared bipolar inputs using spike-triggered independent component analysis (STICA; Fig. 4B) [43, 63]. ST-ICA assumes that the spike-triggered ensemble can be modeled as the product of an unknown mixing matrix and the source signals– here, the upstream input. The resulting outputs are spatial sub-units, whose activations we compute with the checker-board stimulus. We classify subunits as sharing input if their centers are in the same square of the checkerboard and the correlation between the activations is at least 0.5 (Fig. 4C). Using a similar method, subunits have been experimentally shown to map to bipolar cell inputs [64]. To test how meta neuron input affects output information, we compare meta neurons constructed from RGCs with shared bipolar inputs to randomly constructed meta neurons. We find that the distribution of predictive information carried by groups of cells who share at least one subunit is significantly greater than the information carried by other groups (Mann-Whitney U test). Moreover, this difference is maintained after coarse-graining using PCG, CCG, and WTA (Fig. 4D) and in different movies and at different prediction timescales (Fig. 4E). This means that the meta neurons carrying high information in Fig. 2D-E are the ones with convergent inputs, and thus are more representative of the amount of information carried by real downstream neurons than the randomly sampled meta neurons (whose predictive information is clustered near zero). This suggests that the brain has to coarse-grain RGCs together somewhat “carefully” (i.e. not randomly) to preserve predictive information, but previous work in our lab shows a simple way to do this: a downstream cell can learn a PCG compression of predictive information from correlations in its inputs via STDP rules [48]. Therefore, neurons that are correlated by common input might get grouped together downstream to efficiently encode predictive information.

## DISCUSSION

Generating successful behavior in response to changing sensory inputs in the face of sensory and motor delays requires the visual system to make predictions about the future state of the world. Previous work has shown that predictive computations start at the earliest stages of visual processing, in the retina [15, 21–24]. In order for an organism to make use of the predictive information available in the retina, downstream areas must read it out. However, RGCs, the output cells of the retina, show convergence onto many of their downstream areas, limiting the amount of information that can be transferred.

Efficient coding and information bottleneck principles suggest that the near-optimal representation of predictive information available in RGCs [15] is efficiently compressed for downstream readout. Clearly, a coarsegraining function optimized for prediction can achieve this, but it is not yet known whether it is necessary. We compare optimal and prediction-agnostic coarse-grainings of RGC activity to determine whether predictive information can be retained even under agnostic compression. Similarly to previous work in hippocampus and cortex [9, 12], we find that computation-agnostic compression retains substantial information, but we elaborate on previous work by specifying that the information retained is actually about the relevant computation (prediction), and by quantifying how much information is lost by not actively optimizing. We show that in this case, computation-agnostic compression is not overwhelmingly less than optimal. However, efficient coding of predictive information should ideally hold over all natural scenes. We find that an optimal coarse-graining for prediction generalizes across natural scenes, while the prediction-agnostic coarse-grainings do not. Taken together, these results suggest that an optimal coarse-graining is most efficient because it can extract generalizable predictive features. The RGCs may carry so much predictive information that small tweaks to the encoding may still preserve a significant fraction of their predictive power. This kind of shallowness of the predictive information landscape would allow for resilience to wiring damage, or to evolutionary tweaks that slowly move the system towards new and expanded computational goals. Sophisticated machine learning algorithms implemented by deep networks are needed for good generalization across tasks [65–67], so it is striking that a single-layer perceptron can generalize so well in this context. Further work on the origin of this ability to generalize, as in other work from our lab [60], could help make progress on the design of better transfer learning algorithms.

We also explore how RGC input composition affects coarse-grained outputs and show that coarse-graining biologically reasonable inputs yields substantially more predictive information than randomly combined inputs. In the coarse-graining that we use, where optimal readouts are learnable via STDP rules [48], this indicates that synaptic weights could be tuned such that cells with convergent inputs are synaptically summed for near-optimal prediction in all natural scenes.

In the amphibian visual system, the OT is regarded as the main center for visuomotor behavior and object recognition [68, 69], and has evolutionary pressure to use predictive information to make fast motor responses to visual inputs [29–32]. Since we consider our meta neurons as corresponding to OT neurons, our results could be experimentally tested in salamander. This would require tracing RGC inputs to OT neurons, then recording from single neurons in OT to determine how much information they carry about the future of their inputs. This could be compared to the amount of future information carried by different coarse-grainings, to possibly rule out coarse-grainings that do not preserve a comparable amount. The caveat to this is that recording from salamander OT is difficult and not often done, but our results could instead be experimentally tested across species. We do not think what we found is specific to salamanders; our qualitative results about different coarse-grainings should hold in mouse superior colliculus as well, which is involved in making fast defensive and prey behaviors in response to visual inputs [30].

Another compelling future research direction is to use this framework to study different computations and even different sensory systems that exhibit compression. In the retina, prediction is clearly not the only relevant computation [70, 71]; scene identity recognition is especially important for species who travel through significantly different environments, such as salamanders. It would be interesting to optimize PCG on different combinations of prediction and scene identity information to investigate whether it is possible to represent multiple kinds of compressed information near-optimally. This suggests that PCG may be a powerful tool for studying biologically relevant computations under compression.

## MATERIALS AND METHODS

### Multielectrode recordings

Voltage traces from the RGC layer of a larval tiger salamander retina were recorded following the methods described in [72]. Briefly, the retina from a larval tiger salamander was isolated in darkness and a 252 channel multielectrode array was used to record from the retina as images were projected onto the photoreceptor layer. Spikes were sorted using a mostly automated spike sorting algorithm, binned at 60 Hz. This technique captured a highly overlapping neural population of 93 cells that fully covered a region of visual space. The stimuli were 20s movies of trees, grasses, water, fish, and optic flow, which were repeated 83, 80, 84, 91, and 85 times, respectively. We omitted the water movie from our analysis because responses to this movie were extremely sparse. The movies were played in a pseudo-random order, and all natural scenes except for the tree stimulus were displayed at 60 Hz. The tree stimulus was shown at 30 Hz to match the slower recording speed of that camera, and to facilitate comparison with previous recordings in the retina in response to this stimulus [15, 73, 74]. A 30 Hz white noise checkerboard stimulus was played for 30 minutes prior to and after the natural scene stimuli. This dataset is publicly available from Dryad [46].

### Information calculations

Full predictive information is the mutual information between the binary activity pattern 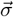 at time t and time *t* + Δ*t*,

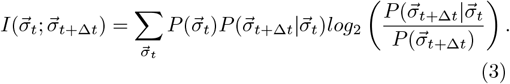

Regardless of the coarse-graining function used to create a meta neuron, predictive information carried by the meta neuron was calculated as the mutual information shared by the meta neuron’s activity and its future input activity: 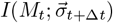. Because we used a small input population size and *M*_*t*_ and 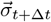 are conditionally independent given 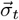, we calculated information from Eq. 1, 2 instead of sampling meta neuron activity to estimate the entropies. To find the error in the mutual information estimates, we followed the methods in [75]: data were subsampled via bootstrapping for 50% of the data, with 50 bootstrap samples taken. Errors were estimated as the standard deviation of the sampled information, divided by 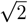.

### Coarse-graining functions

Our coarse-graining functions compressed *N* cells into one meta neuron by transforming a binary activity vector with 2^*N*^ states into an activity vector with only 2 states. The output of PCG was the conditional probability that the meta neuron spiked given a certain pattern of current binary activity, which was modeled as a weighted sum of the inputs passed through a sigmoid nonlinearity

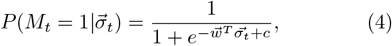

where 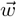 represents the weights and *c* represents the threshold parameter of the nonlinearity. Since meta neurons only have two states, we have fully characterized 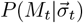. The other components of Eq. 2, 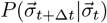 and 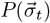, were counted directly from the data.

To construct a CCG meta neuron, inputs were summed together based on their pairwise correlation: the first two maximally correlated cells were summed together, then the next two maximally cells were summed, and so on. These were normalized by their maximal values, and then the process was repeated until there was only one trace left. Note that the process could be stopped earlier to obtain a population of meta neurons. The trace was bounded between 0 and 1, and we took it to represent the spiking probability of the meta neuron. We used this to simulate the meta neuron’s activity, and for each input pattern 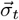, we counted the number of times the meta neuron elicited an output spike to calculate 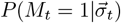.

The output of WTA was simply the trace of the input cell with the highest firing rate. Again, we counted the number of times the meta neuron spiked in response to each input pattern 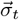 to calculate 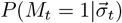.

### Optimization Methods

We maximized Eq. 1 for PCG over the weights and threshold parameter, subject to the constraint that its firing probability was equal to the CCG meta neuron with the same input cells. The firing probability of PCG is

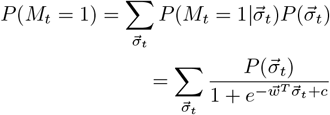

which makes clear how its firing probability depends on the weights and threshold parameter. To perform constrained optimization, we used the `fmincon` function in MATLAB. We initialized the parameters with 100 random seeds, bounded the weights between [0, 100], and bounded the threshold between [0, 200]. We did not specify the gradient of the objective function `(‘GradObj’ = off)` and we used the Sequential Quadratic Programming (SQP) optimization algorithm.

Random PCG meta neurons were constructed for each movie by randomly drawing weights from the distribution of optimal PCG weights found in the relevant natural movie, then adjusting the sigmoid threshold parameter so that its firing probability matched the firing probability of the corresponding optimal PCG meta neuron with the same inputs. When testing a PCG meta neuron on a different movie from which it was trained, we used the training weights but adjusted the threshold parameter so that the firing rate probability matched that of the corresponding PCG meta neuron optimized on the test movie.

### Spike-Triggered ICA

Spike-Triggered ICA (ST-ICA) was calculated following methods developed in [43]. Briefly, we computed the spike triggered average for each cell, then decomposed the resulting spatiotemporal filter into spatial and temporal filters. We then convolved the checkerboard stimulus with the temporal filter before applying ICA. The resulting output sources were spatial subunits, which we identified as the same following methods developed in [76].

### Drawing input cells

73 input cell sets of size *N* = 4 were drawn from the population of 93 total recorded cells for Figs. 2-4. These were a combination of random cells, non-overlapping sets made by sorting and grouping cells by their individual predictive information, and subsamples of the top 20 most individually predictive cells. The “shared input” sets (*n* = 19) in Fig. 3 were classified as the sets where at least 2 of the 4 input cells shared at least 1 ST-ICA spatial subunit. Difference in the distributions between shared input sets and other sets was assessed with a Mann Whitney U test: for each combination of coarse-graining (full, PCG, CCG, and WTA), movies, and prediction horizons (Δ*t* = 16 ms and Δ*t* = 100 ms), we used the median as the measure of central tendencies of the shared input and other groups. We performed the U test with the “ranksum” function in MATLAB. For each combination, the distributions in the shared input and other groups differed significantly (*P <* .001 two-tailed).

### Data, Materials, and Software Availability

The data have been published at https://doi.org/10.5061/dryad.4qrfj6qm8 [46].

## ACKNOWLEDGMENTS

This work was supported in part by the National Science Foundation (NSF), through the Physics Frontier Center for Living Systems (PHY-2317138) and a Graduate Research Fellowship Grant (DGE-214000) (SCLD); and by the NSF-Simons National Institute for Theory and Mathematics in Biology, (NSF DMS-2235451 and Simons Foundation MP-TMPS-00005320).

## Notes

https://doi.org/10.5061/dryad.4qrfj6qm8

### Competing Interest Statement

The authors have declared no competing interest.

