## Supplemental Appendix 1 for "Preserving predictive information under biologically plausible compression"

### Supporting Information Appendix (SI)

#### Predictive PCG generalizes for longer timescales

We repeat our generalization analysis for a longer prediction horizon, by training and testing PCG meta neurons on different individual movies. We choose a prediction horizon of $\Delta t = 100$ ms because by then, the autocorrelations in the checkerboard movie have died off, while there are still nonzero autocorrelations in the natural movies (Fig. S1A). Therefore, at $\Delta t = 100$ ms, the predictive information in the meta neurons is entirely stimulus-driven. As expected, we find nonzero predictive information in PCG meta neurons trained on the natural movies and close to zero information in PCGs trained on the checkerboard (Fig. S1B). Generalization across movies is slightly worse than for $\Delta t = 16$ ms, but is still notable (Fig. S1C-D).

#### Predictive PCG generalizes over timescales

We investigate whether PCG generalizes over prediction horizons as well as natural scenes by training and testing PCG meta neurons on different timescales. Indeed, PCG maintains close to optimal future information when tested on a different prediction timescale from which it was trained (Fig. S2A). Moreover, future information maintained over a different training horizon and training movie is comparable in different test movies (Fig. S2B-D).

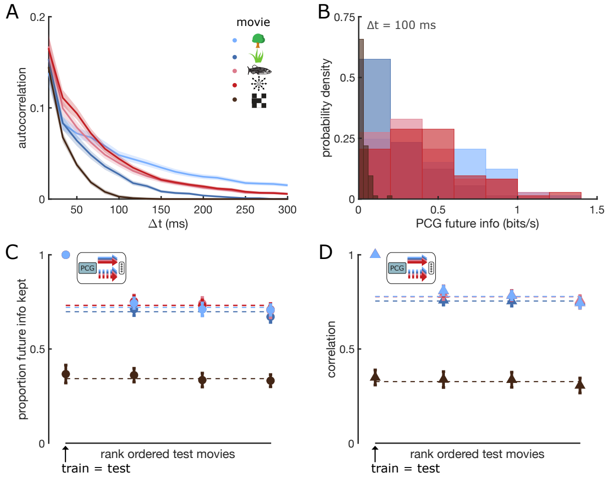


FIG. S1. **PCG generalizes for longer timescales.** A) Autocorrelation timescales for each movie. B) Distributions of future information in PCG meta neurons in the different movies. C) Mean proportion future ($\Delta t = 100$ ms) information retained by PCG trained and tested on different movies. D) Mean correlation coefficients between PCGs trained on different movies.

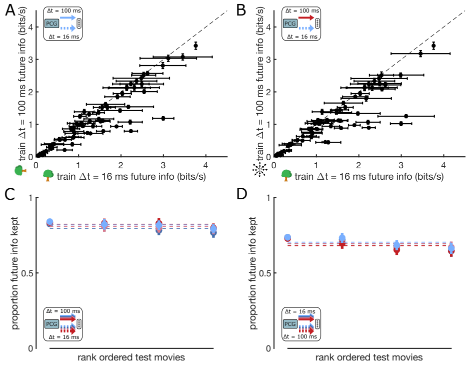


FIG. S2. **PCG generalizes across timescales.** A) Future information maintained by PCGs trained on predicting $\Delta t = 100$ ms in the future on the tree movie, then tested on prediction $\Delta t = 16$ ms in the tree movie. B) Future information maintained by PCGs trained on predicting $\Delta t = 100$ ms in the future on the tree movie, then tested on prediction $\Delta t = 16$ ms in another movie (optic flow). C) Mean proportion future information retained by PCG trained on prediction $\Delta t = 100$ ms in the future and tested on prediction $\Delta t = 16$ ms in the future in different movies. D) Mean proportion future information retained by PCG trained on prediction $\Delta t = 16$ ms in the future and tested on prediction $\Delta t = 100$ ms in the future in different movies.
